# Intranasal Immunization with a Proteosome-Adjuvanted SARS-CoV2 Spike Protein-Based Vaccine is Immunogenic and Efficacious in Mice & Hamsters

**DOI:** 10.1101/2022.03.02.482651

**Authors:** Felicity C. Stark, Bassel Akache, Lise Deschatelets, Anh Tran, Matthew Stuible, Yves Durocher, Michael J. McCluskie, Gerard Agbayani, Renu Dudani, Blair A. Harrison, Tyler M. Renner, Shawn R. Makinen, Jegarubee Bavananthasivam, Diana Duque, Martin Gagne, Joseph Zimmermann, C. David Zarley, Terrence R. Cochrane, Martin Handfield

## Abstract

With the persistence of the SARS-CoV-2 pandemic and the emergence of novel variants, the development of novel vaccine formulations with enhanced immunogenicity profiles could help reduce disease burden in the future. Intranasally delivered vaccines offer a new modality to prevent SARS-CoV-2 infections through the induction of protective immune responses at the mucosal surface where viral entry occurs. Herein, we evaluated a novel protein subunit vaccine formulation containing a resistin-trimerized prefusion Spike antigen (SmT1v3) and a proteosome-based mucosal adjuvant (BDX301) formulated to enable intranasal immunization. In mice, the formulation induced robust antigen-specific IgG and IgA titers, in the blood and lungs, respectively. In addition, the formulations were highly efficacious in a hamster challenge model, reducing viral load and body weight loss. In both models, the serum antibodies had strong neutralizing activity, preventing the cellular binding of the viral Spike protein based on the ancestral reference strain, the Beta (B.1.351) and Delta (B.1.617.2) variants of concern. As such, this intranasal vaccine formulation warrants further development as a novel SARS-CoV-2 vaccine.

## Introduction

The etiological agent of the COVID-19 pandemic, SARS-CoV-2, has proven to be highly virulent and adaptable. Despite the development and deployment of multiple relatively safe and efficacious vaccines worldwide, the emergence of novel viral variants with decreased sensitivity to vaccine-induced immune responses has caused the pandemic to persist and still present a major challenge to human health ^1–3^.

In the initial response to the COVID-19 pandemic, vaccine developers were able to produce efficacious vaccines targeting SARS-CoV-2 with unprecedented speed by utilizing several established and novel vaccine platforms. Inactivated virus vaccines (e.g. Sinopharm’s BBIBP-CorV and Sinovac’s CoronaVac), as well as mRNA- (e.g. Pfizer-BioNTech’s Comirnaty® and Moderna’s Spikevax®) and viral vector-based vaccines (e.g. Oxford-AstraZeneca’s Vaxzevria® and Janssen/Johnson & Johnson’s Ad26.COV2.S) quickly advanced through clinical trials to regulatory approval due to their strong immunogenicity/efficacy, but also in part to their relatively rapid manufacturing processes ^4–6^. As such, they were the first wave of SARS-CoV-2 vaccines to become widely available in many parts of the world including China, North America and Europe. However, questions remain regarding their efficacy and safety in the long term. Although rare in frequency, severe cases of anaphylaxis, myocarditis and/or thrombocytopenia have been linked specifically to the mRNA and viral vector vaccine platforms^7–9^. With the emergence of novel COVID-19 variants of concern capable of partially evading the protection induced by the currently approved vaccines ^1–3,10^, the development of novel vaccines with improved safety and efficacy profiles could reduce the impact of SARS-CoV-2 and its variants in the future.

SARS-CoV-2 is an airborne pathogen that enters the body primarily through the upper respiratory tract (URT; nose and mouth). The URT is likely the initial location of infectivity as it contains a much greater concentration of ACE2 receptors than the lower respiratory tract ^11**Error! Reference source not found**.^. Immune responses initiated in these mucosal sites often involve mucosa-associated lymphoid tissues (MALT), which are a compartmentalized immunological system that functions independently from the systemic immune apparatus^12^. This localized system includes epithelial cells that take up antigen, which is subsequently shuttled to or captured by antigen presenting cells (APCs, i.e. dendritic cells, macrophages) and presented to CD4^+^/CD8^+^ T-cells all located within the mucosal inductive site^13^. Intranasal vaccines have the potential to induce systemic immune responses as with parenteral vaccines, but can also generate robust broadly protective mucosal immune responses based on secretory IgA antibodies and tissue-resident T cells at the mucosal surface of the respiratory tract at the site of viral entry potentially limiting infection before it becomes established^14^. This in turn could lead to reduced transmission of SARS-CoV-2 in vaccinated populations relative to vaccines delivered parenterally. In addition, due to the non-invasive nature of vaccine administration, it may increase vaccine uptake in the wider population.

Mucosal adjuvants are considered essential in activating mucosal immunity to overcome the natural tolerance to antigens encountered in this compartment^15^. A number of adjuvants have demonstrated an ability to potentiate adaptive and innate immunity when administered on a mucosal surface^16–18^. BDX301 is a Proteosome-based adjuvant composed of porin proteins and lipooligosaccharides from *Neisseria meningitidis*. Proteosomes are a family of mucosal adjuvants derived from the outer membranes of Gram-negative pathogenic bacteria and have shown strong activity when delivered intranasally, inducing high levels of antigen-specific IgA within the respiratory tract^19–21^. In fact, influenza vaccines containing these types of adjuvants have been shown to be safe and effective in human clinical trials^22,23^. Previously, a nasal vaccine comprised of a SARS-CoV-1 recombinant Spike protein with another Proteosome based adjuvant, Protollin, elicited similar serum neutralizing antibody levels as an aluminum hydroxide-adjuvanted formulation containing the same antigen administered intramuscularly, but had significantly lower and undetectable lung SARS-CoV-1 viral titres at 3 days post-challenge, compared to aluminum hydroxide, in a mouse model^24^.

With the persistence of SARS-CoV-2, vaccines based on different technologies and routes of administration warrant investigation. We have previously demonstrated the strong immunogenicity of a resistin-trimerized SARS-CoV-2 Spike antigen, referred to as SmT1^25^. Intramuscular administration of adjuvanted vaccine formulations of SmT1 induced robust immune responses that were strongly protective in a hamster challenge model. Herein, we evaluated the immunogenicity and efficacy of intranasal vaccine formulations based on a FLAG and histidine tag-less version of SmT1 (i.e. SmT1v3) and the mucosal adjuvant BDX301 in preclinical models. Overall, we demonstrate that intranasal administration of the vaccine formulations induces high levels of antigen-specific IgG and IgA in mice and is efficacious in a hamster challenge model, preventing body weight loss and eliminating viral load in lungs/nasal turbinates.

## Results

### IgG antibody response in immunized mice

BALB/c mice (n=10/group) were immunized by intranasal instillation with vaccine formulations consisting of SmT1v3 alone or adjuvanted with BDX301 on Days 0 and 21. SmT1v3 based on the ancestral reference strain originally identified in Wuhan was used for all vaccine formulations in this study. Controls included mice injected with the vaccine vehicle, phosphate-buffered saline (PBS), or SmT1v3 adjuvanted with aluminum phosphate (AdjuPhos^™^). Aluminum phosphate is a conventional vaccine adjuvant found in a number of approved vaccines and is a potent inducer of antigen-specific antibody responses^26,27^. SmT1v3-AdjuPhos^™^ formulations were delivered intramuscularly^28^. To assess the ability of the vaccine formulations to induce systemic immune responses following one or two vaccine doses, serum samples were taken on Days 21 and 35, respectively.

Following a single vaccine dose, SmT1v3 adjuvanted with BDX301 or aluminum phosphate induced statistically similar anti-Spike IgG Geomean Titers (GMT) of 5,288 and 7,091 (p>0.05), respectively, in the sera of immunized mice (Fig. 1A). These levels were significantly greater than those observed in animals immunized with antigen alone or vehicle control (p<0.0001). Antigen-specific antibody titers were increased >20-fold in the sera of mice following administration of a second vaccine dose of SmT1v3 adjuvanted with BDX301 or aluminum phosphate, with measured GMTs of 392,690 and 163,003, respectively (p>0.05; Fig. 1B). Again, they were significantly greater than both control groups (p<0.0001), which still showed low or non-detectable antibody titers.

**Figure 1:**
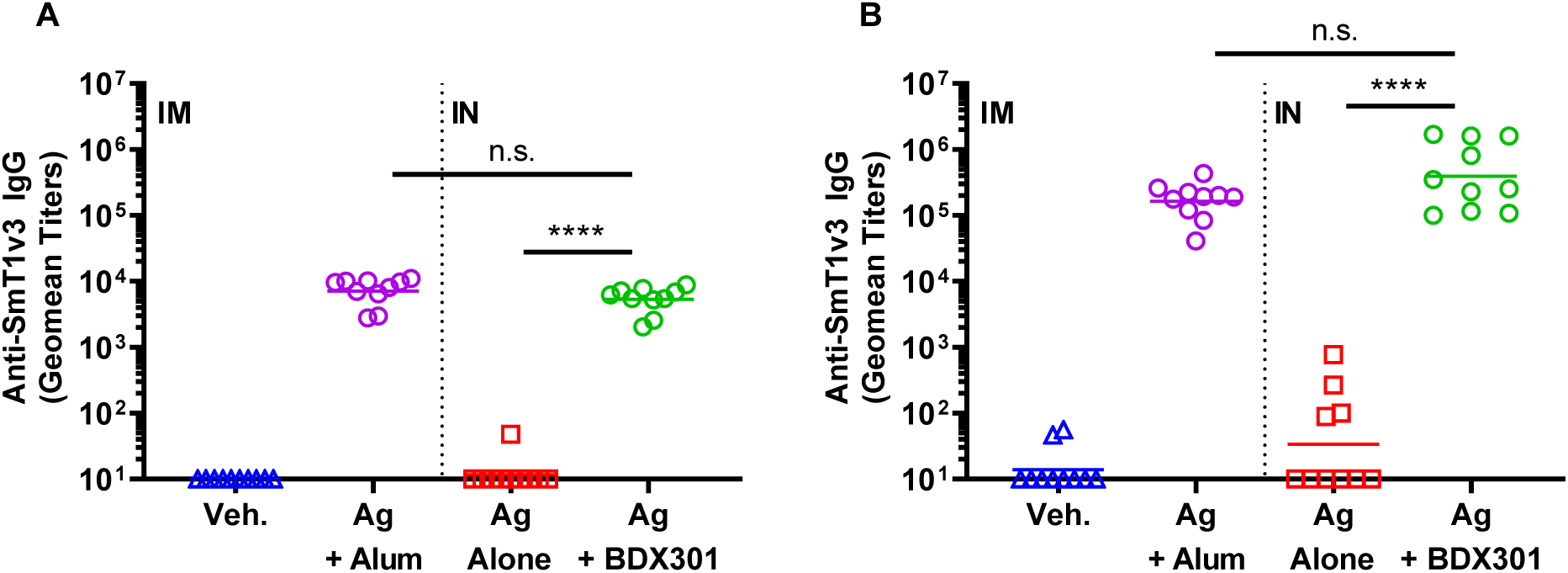
Anti-Spike IgG titers induced by SmT1v3 and BDX301 formulations in mice. BALB/c mice (n=10/group) were immunized twice on Days 0 and 21. Vehicle control (Veh.) or the antigen (Ag) SmT1v3 (10 µg) with aluminum phosphate (Alum) (100 µg) were administered via the intramuscular (IM) route, while SmT1v3 (10 µg) with or without BDX301 (5 µg) were administered via the intranasal (IN) route. Serum collected 21 days after the 1^st^ dose (Panel A) and 14 days after the 2^nd^ dose (Panel B) were analyzed by ELISA to determine the levels of antigen-specific IgG titers. Antibody titers are expressed as a reciprocal value of the serum dilution calculated to generate an OD450 = 0.2. For statistical analysis, antibody titers were log-transformed and then analyzed by a one-way ANOVA with Tukey’s multiple comparisons test. ****: p<0.0001, n.s.: no significant difference.

### Neutralization response in immunized mice

While the induction of anti-Spike IgG antibodies is currently thought to be a strong predictor of vaccine efficacy against COVID-19^29^, the functionality of these antibodies as measured by their ability to prevent viral Spike binding to the cells and prevent infection is especially important. Using a surrogate cell-based neutralization assay previously shown to have a strong correlation to responses obtained with viral-based SARS-CoV-2 assays ^28^, we evaluated the neutralization activity of the immunized mouse serum against the ancestral reference strain and Beta variant of concern (VOC). The assay measures the ability of serum antibodies to prevent binding of Spike protein to the ACE2 receptor on the surface of VERO E6 cells. Mutations within the Spike protein from the Beta VOC, have made it especially resistant to neutralization by antibodies induced by the reference strain ^30^. SmT1v3-BDX301 vaccine formulations induced strong neutralizing activity against both the ancestral and Beta SARS-CoV2 Spike proteins, which was significantly higher than with antigen alone (Fig. 2; p<0.0001). Additionally, the BDX301-adjuvanted formulation induced significantly greater neutralization than that observed with the SmT1v3-aluminum phosphate formulation against the Beta variant (71 vs. 35%; p<0.05), while neutralization was statistically similar against the Spike from the ancestral reference strain (93 vs. 80%; p>0.05).

**Figure 2:**
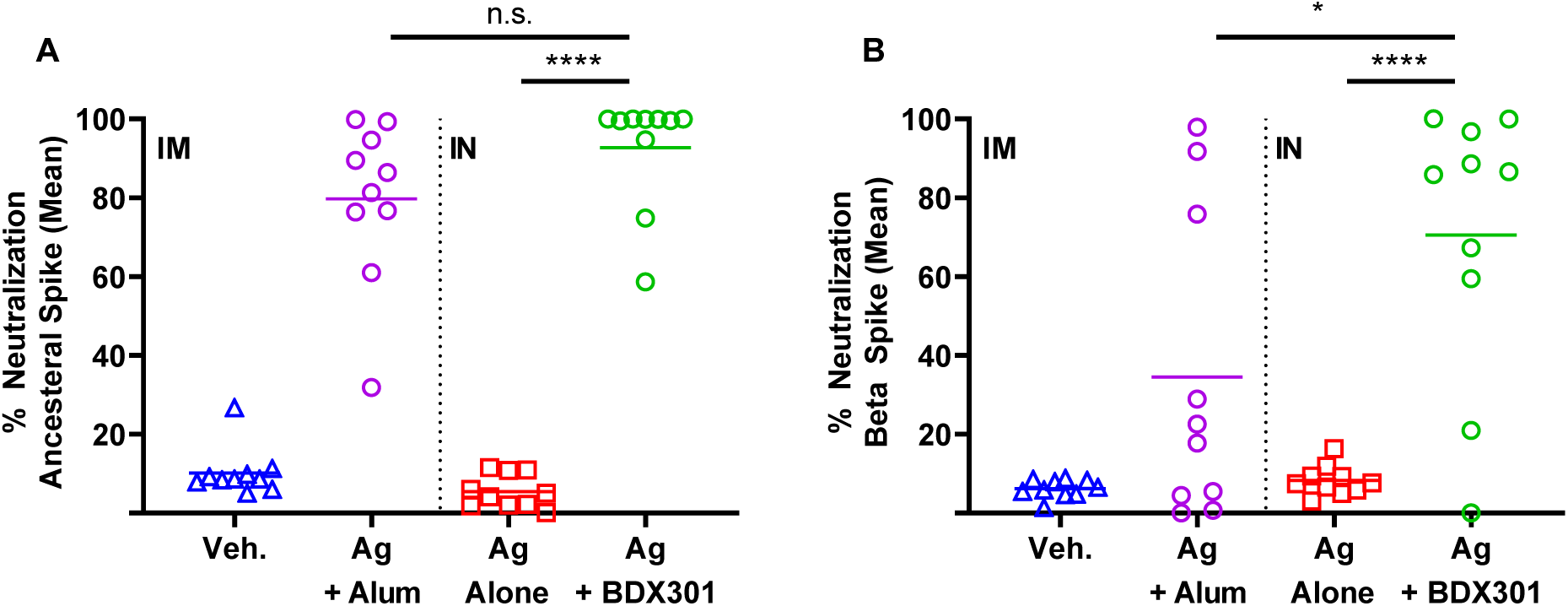
Neutralization activity induced by SmT1v3 and BDX301 vaccine formulations in mice. BALB/c mice (n=10/group) were immunized twice on Days 0 and 21 with PBS (vehicle control, Veh.) or the antigen (Ag) SmT1v3 (10 µg), with or without aluminum phosphate (Alum) (100 µg), via the intramuscular (IM) route or SmT1v3 (10 µg), with or without BDX301 (5 µg), via the intranasal (IN) route. Serum collected on Day 35 was diluted 75-fold and analyzed for its ability to block binding of Spike protein based on the ancestral reference strain (Panel A) or Beta variant (Panel B) to VERO E6 cells. The neutralization activity is calculated as a percent reduction from signal seen with control cells incubated in the absence of serum. For statistical analysis, values were analyzed by a one-way ANOVA with Tukey’s multiple comparisons test. ****: p<0.0001, *:p<0.05, n.s.: no significant difference.

### IgA antibody response in immunized mice

As enhancement of local IgA responses in the mucosal compartment is typically one of the main advantages of mucosal proteosome-adjuvanted vaccine formulations, antigen-specific IgA titers were also assessed in the bronchoalveolar lavage (BAL) fluid and serum of mice collected on Day 35. Only SmT1v3-BDX301 immunized mice showed detectable anti-SmT1v3 IgA titers, as titers in mice immunized with antigen alone or SmT1v3-aluminum phosphate were below the assay’s limit of detection (Fig. 3). In both the BAL fluid and serum, IgA titers were significantly higher than the control groups with a GMT of 1,000 and 5,074 in the BAL and serum, respectively (p<0.0001). Interestingly, the anti-SmT1v3 IgG/IgA ratio was ∼25-fold lower in the BAL vs. serum (p<0.01; Fig. 3D), indicating that the antigen-specific IgA detected in the BAL was not simply due to cross-contamination with blood or leakage of blood through the mucosal epithelium. Having established the immunogenicity of a vaccine formulation based on BDX301 and SmT1v3 in mice, we next sought to evaluate its efficacy against a live SARS-CoV-2 challenge in hamsters.

**Figure 3:**
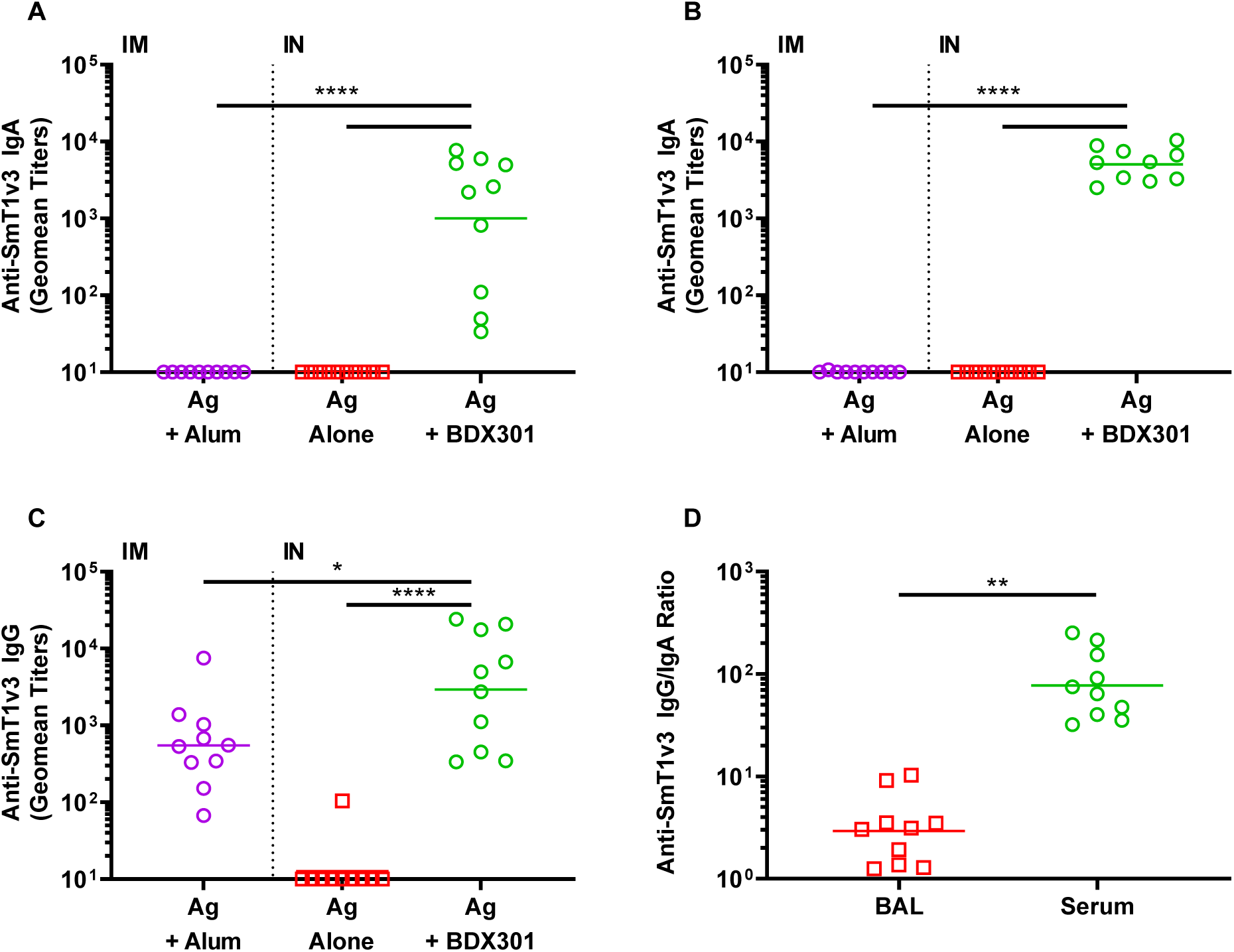
Comparison of anti-Spike IgA and IgG titers induced by SmT1v3 and BDX301 formulations in serum and lungs of mice. BALB/c mice (n=10/group) were immunized twice on Days 0 and 21. The antigen (Ag) SmT1v3 (10 µg) with aluminum phosphate (Alum) (100 µg) was administered via the intramuscular (IM) route, while SmT1v3 (10 µg), with or without BDX301 (5 µg), were administered via the intranasal (IN) route. Bronchoalveolar lavage (BAL; Panel A) or serum (Panel B) collected on Day 35 were analyzed by ELISA to determine the levels of antigen-specific IgA titers. Levels of antigen-specific IgG titers were also measured in the BAL (Panel C) and used along with serum IgG titers from Figure 1B to determine the anti-SmT1v3 IgG/IgA ratios in both BAL and serum for the Ag+BDX301-immunized mice (Panel D). Antibody titers are expressed as a reciprocal value of the BAL or serum dilution calculated to generate an OD450 = 0.2. For statistical analysis, antibody titers were log-transformed and then analyzed by a one-way ANOVA with Tukey’s multiple comparisons test, while IgG/IgA ratios were analyzed by a Student’s t-test. ****: p<0.0001, **:p<0.01, *:p<0.05.

### IgG antibody response in immunized hamsters

Golden Syrian hamsters are a widely accepted model to evaluate vaccines against SARS-CoV-2, due to their susceptibility to infection as well as the manifestation of similar disease pathology in the lungs to that seen in COVID-19 patients with pneumonia^31,32^. Golden Syrian hamsters were immunized intranasally twice on Days 0 and 21 with BDX301 alone, or in combination with 5 or 15 µg of SmT1v3. A negative control group dosed with PBS alone was also included. We first sought to measure the immunogenicity of the vaccine formulations in hamsters. Serum samples were collected on Days 20 and 35 to assess anti-SmT1v3 IgG titers. As expected, both doses of SmT1v3-BDX301 induced high levels of antigen-specific IgG titers that were significantly higher than seen in the antigen alone or vehicle control groups at either timepoint (Fig. 4; p<0.0001). At Day 20, significantly greater GMTs of anti-SmT1v3 IgG were induced by BDX301 formulations containing 15 µg vs. 5 µg antigen (2,766 vs. 718; p<0.001; Fig. 4A). By Day 35, 14 days post the second immunization, the GMT titers were still higher with 15 µg vs. 5 µg, but they did not reach a level of statistical significance (19,492 vs. 8,510; p>0.05; Fig. 4B).

**Figure 4:**
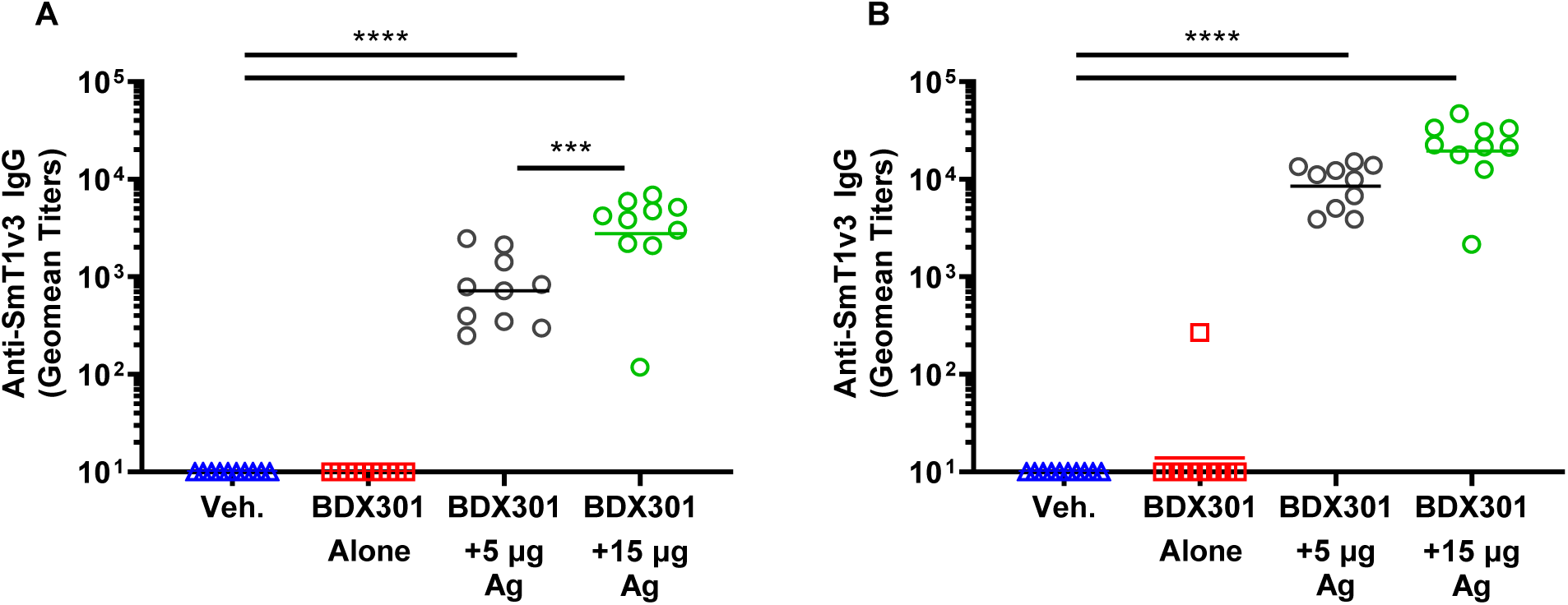
Anti-Spike IgG titers induced by SmT1v3 and BDX301 formulations in hamsters. Syrian Golden hamsters (n=10/group) were immunized twice on Days 0 and 21 with PBS (vehicle control, Veh.) delivered intramuscularly or BDX301 (5 µg) with or without SmT1v3 (5 µg or 15 µg) via the intranasal route. Serum collected on Day 20 (Panel A) and Day 35 (Panel B) were analyzed by ELISA to determine the levels of antigen-specific IgG titers. Antibody titers are expressed as a reciprocal value of the serum dilution calculated to generate an OD450 = 0.2. For statistical analysis, antibody titers were log-transformed and then analyzed by a one-way ANOVA with Tukey’s multiple comparisons test. ***: p<0.001, ****: p<0.0001.

### Neutralization response in immunized hamsters

As with mice, the neutralization activity of the immunized hamster serum was assessed, but in addition to the ancestral reference strain and Beta VOC, the Spike protein from the Delta VOC was also included. The BDX301 formulations with either 5 µg or 15 µg antigen induced strong neutralization activity to the 3 different Spike variants. The strongest neutralization activity was seen against the ancestral reference strain, with BDX301 with 5 or 15 µg of SmT1v3 inducing an average of 55 and 71% neutralization, respectively (p<0.0001 vs. vehicle or adjuvant alone groups; Fig. 5A). While neutralization activity against the Beta variant was detected in many animals immunized with BDX301-SmT1v3, it did not reach a level of statistical significance when compared to the control groups (Fig. 5B). Neutralization activity against the Delta variant was quite strong, with an average of 39 and 52% neutralization measured in the sera of animals immunized with BDX301 with 5 and 15 µg of SmT1v3, respectively. (p<0.05 vs. control groups; Fig. 5C). A plaque reduction neutralization test was also conducted to test the ability of the serum samples to block infection of VERO E6 cells by live SARS-CoV-2 (ancestral reference strain). Either antigen dose combined with BDX301 induced significantly greater neutralization compared to vehicle or adjuvant alone (p<0.0001) (Fig. 5D).

**Figure 5:**
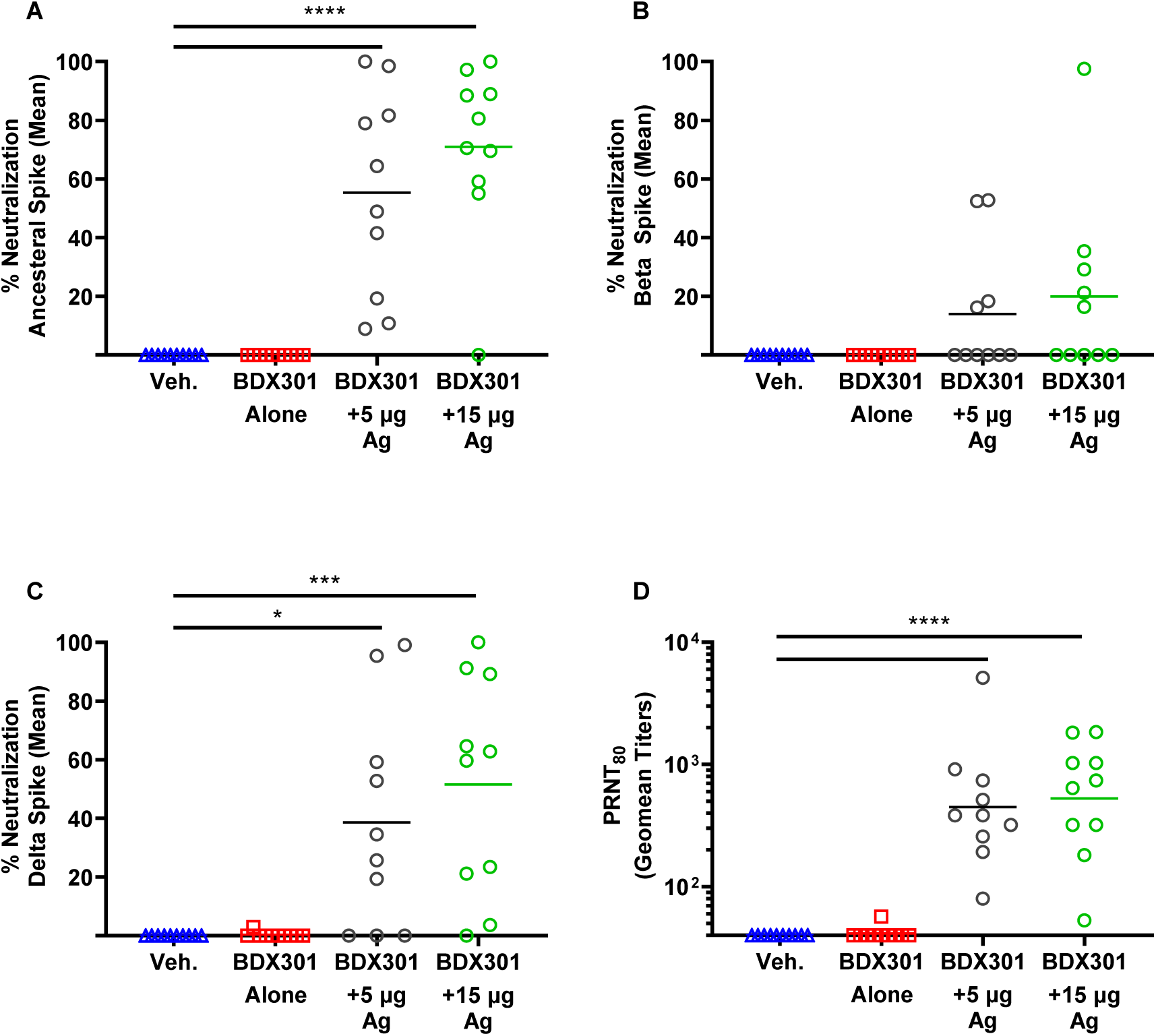
Neutralization activity induced by SmT1v3 and BDX301 formulations in hamsters. Syrian Golden hamsters (n=10/group) were immunized twice on Days 0 and 21 with PBS (vehicle control, Veh.) delivered intramuscularly or BDX301 (5 µg) with or without SmT1v3 (5 µg or 15 µg) via the intranasal route. Serum collected on Day 35 was diluted 25-fold and analyzed for its ability to block binding of Spike protein based on the ancestral reference strain (Panel A), Beta variant (Panel B) or Delta variant (Panel C) to VERO E6 cells. The neutralization activity is calculated as a percent reduction from signal seen with control cells incubated in the absence of serum. Sera from Day 35 was also tested by a plaque reduction neutralization test for its ability to block infection of VERO cells by SARS-CoV-2 (ancestral reference strain). PRNT titers are expressed as a reciprocal value of the serum dilution calculated to generate an 80% reduction from the number of plaques observed in control wells without serum. For statistical analysis, values were analyzed by a one-way ANOVA with Tukey’s multiple comparisons test. ****: p<0.0001, ***: p<0.001, *:p<0.05, n.s.: no significant difference.

### SARS-CoV-2 viral challenge of immunized hamsters

On Day 42, hamsters were challenged with 1 × 10^5^ PFU of SARS-CoV-2 (ancestral reference strain) and monitored daily for body weight change post-challenge. At 5 days post-challenge, the BDX301 adjuvant alone and PBS vehicle control groups lost ∼9 and 11% of their body weight, respectively, with no significant difference between the two (Fig. 6). Hamsters immunized with either 5 or 15 µg of SmT1v3 were protected from body weight loss over the course of the study, demonstrating significantly less body weight loss than both control groups (p<0.0001; Fig.6A). On Day 47, hamsters were euthanized and viral titers were quantified in lung and nasal turbinates by plaque assay. While viral load was high in control animals immunized with PBS or adjuvant alone, no viral titers were detected in tissues of animals immunized with BDX301 combined with 5 or 15 µg of SmT1v3 (Fig. 6B & 6C). Importantly, as the adjuvant alone group did not protect hamsters from body weight loss or viral replication, we can deduce that the observed protection in this study was largely mediated by antigen-specific responses and not innate immune activation.

**Figure 6:**
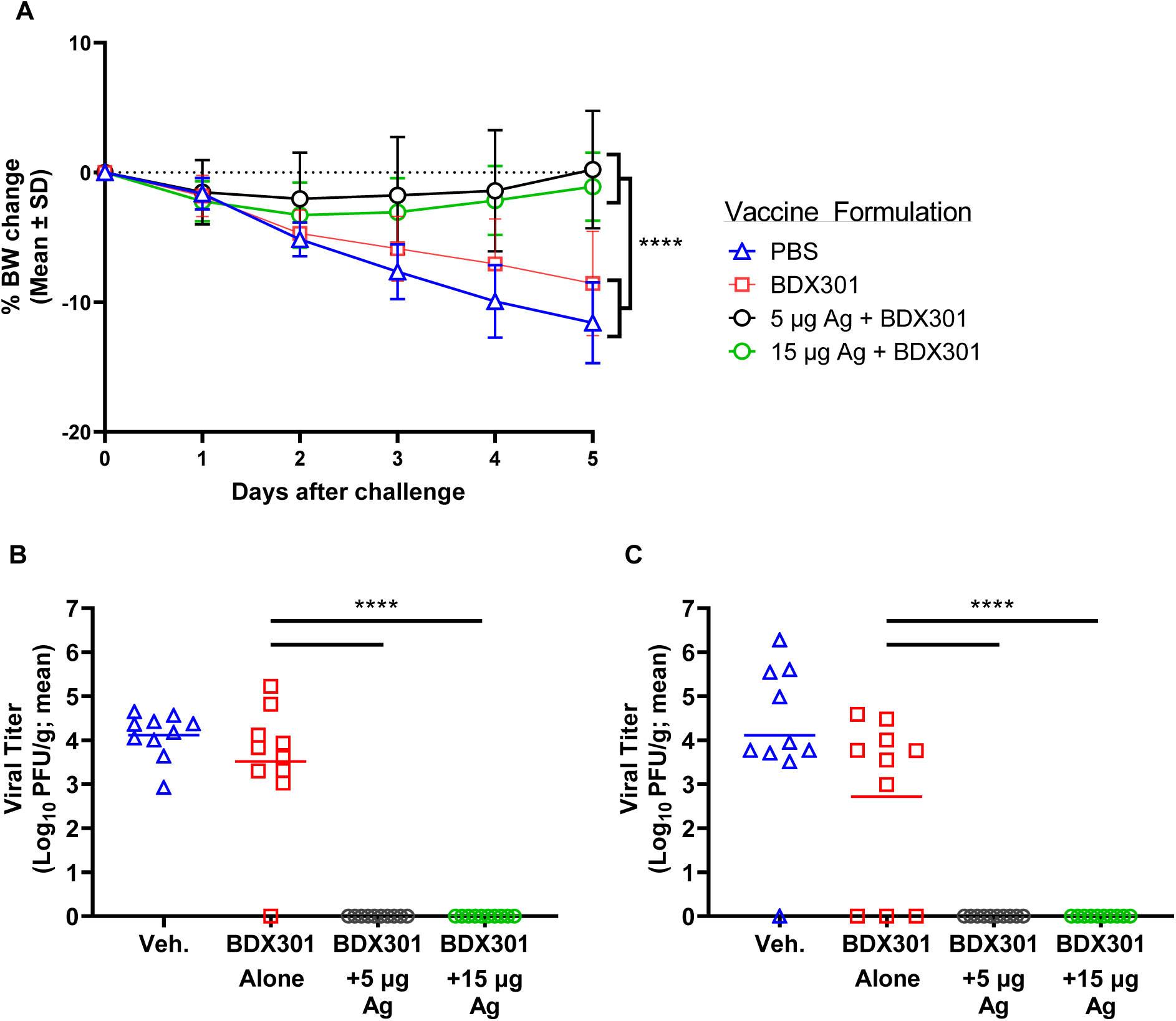
Efficacy of SmT1v3 and BDX301 formulations against SARS-CoV-2 viral challenge in hamsters. Syrian Golden hamsters were immunized twice on Days 0 and 21 with PBS (vehicle control, Veh.) delivered intramuscularly or BDX301 (5 µg) with or without SmT1v3 (5 or 15 µg) via the intranasal route. On Day 42 all hamsters were challenged with 1 × 10^5^ PFU of SARS-CoV-2. Hamsters were monitored daily for body weight change post-challenge (Panel A). On Day 47, hamsters were euthanized and viral titers were quantified in lung (Panel B) and nasal turbinates (Panel C) by plaque assay. For statistical analysis, a two-way (Panel A) or one-way (Panels B &C) ANOVA with Tukey’s multiple comparisons test was performed. ****: p<0.0001, ***: p<0.001.

## Discussion

The purpose of this study was to first conduct an immunogenicity comparison of the intranasal adjuvant BDX301 using the conventional intramuscular adjuvant aluminum phosphate (AdjuPhos^™^) as a benchmark in mice with the ancestral Wuhan strain recombinant Spike protein SmT1v3, a FLAG and histidine tag-less version of SmT1, which was previously described elsewhere^25^. In the second part of our study, we assessed the inhibition of virus replication in hamsters immunized with SmT1v3 formulated with BDX301, hereafter referred to as NT-CoV2-1. Two intranasal doses of NT-CoV2-1 elicit strong serum IgG binding and neutralization responses to the ancestral Wuhan strain that are comparable to those seen following intramuscular dosing with SmT1v3 formulated with aluminum phosphate. Neutralization responses induced by NT-CoV2-1 to the Beta (B.1.351) variant were significantly higher than observed with the aluminum phosphate-adjuvanted formulation (p<0.05). No measurable responses were seen with the antigen alone controls.

A key differentiating feature of intranasal administration of NT-CoV2-1 is the IgA response seen in BAL and serum in mice, which was not seen in mice following intramuscular administration of SmT1v3 formulated with aluminum phosphate or intranasal immunization with SmT1v3 alone. These results are consistent with other animal studies using intranasally administered recombinant SARS Spike protein^24^ and vector-expressed SARS-CoV-2 Spike protein constructs^33^ and suggest that intranasal immunization may be necessary to induce BAL IgA antibodies.

In hamsters, NT-CoV2-1 induced high levels of antigen-specific IgG titers and neutralization activity to the ancestral Wuhan strain, as well as neutralization activity against the Beta (B.1.351) and Delta (B1.617.2) variants, though at reduced potency than for Wuhan. Interestingly, the Delta variant appeared more susceptible than the Beta variant to neutralization by antisera induced by the ancestral Wuhan strain, as has been reported elsewhere in the literature^34^. The intranasal challenge of hamsters receiving two intranasal doses of NT-CoV2-1 confirmed the functional activity of the immune response, as hamsters maintained weight and showed no measurable virus titer in lungs or nasal turbinates. Control hamsters showed significant weight loss and residual virus in both tissues. These results are consistent with the benchmark set with other COVID-19 vaccine candidates that have proceeded to clinical trial.

Intramuscularly administered COVID-19 vaccines have proven highly effective in preventing COVID-19 symptomatic disease, hospitalization and death. However, recent experience with these vaccines suggest they may not be as effective in preventing virus transmission, particularly with newly emerging variants such as Delta and Omicron ^35^. It has been posited that induction of local immunity in the nasal-pharyngeal cavity by intranasal vaccine administration could reduce viral shedding and transmission, which has been supported by recent studies in hamsters and rhesus macaques^36,37^. Hence there is growing interest in developing an intranasal COVID-19 vaccine with potential to reduce virus transmission more effectively^38^.

## Conclusion

NT-CoV2-1 is composed of a recombinantly expressed ancestral Wuhan Spike protein trimer (SmT1v3) adjuvanted with a proteosome-based intranasal adjuvant BDX301^19–21^. We have demonstrated that NT-CoV2-1 induces superior mucosal and comparable systemic immune responses to aluminum phosphate-adjuvanted formulations in mice and is able to suppress SARS-CoV-2 infection in hamsters. Unlike vector-based intranasal vaccine candidates^39^, protein subunit vaccines such as NT-CoV2-1 can be enhanced with the addition of an adjuvant and avoid the potential inhibition due to anti-vector antibodies induced by prior administration of the same vector. These results support further development of the NT-CoV2-1 vaccine.

## Materials and Methods

### Animals and Virus

Female BALB/c mice (8-12 weeks old) and both male and female golden Syrian hamsters (81-90 g) were obtained from Charles River Laboratories (Saint-Constant, Canada). Animals were maintained at the small animal facility of the National Research Council Canada (NRC) in accordance with the guidelines of the Canadian Council on Animal Care. All procedures performed on animals in this study were in accordance with regulations and guidelines reviewed and approved in animal use protocol 2020.06 & 2020.10 by the NRC Human Health Therapeutics Animal Care Committee.

The ancestral reference strain of SARS-CoV-2 (isolate Canada/ON/VIDO-01/2020 obtained from the National Microbiology Lab, Winnipeg, Canada) was propagated and quantified on Vero E6 cells. Whole viral genome sequencing was carried out to confirm exact genetic identity to original isolate. Passage 3 virus stocks were used in all subsequent experiments.

### Vaccine antigen and adjuvants

Recombinant SmT1v3 Spike trimer constructs are based on “tagless” versions of the SARS-CoV-2 Spike trimers described previously ^40,41^. Briefly, the SARS-CoV-2 reference strain Spike ectodomain sequence (amino acids 1-1208 derived from Genbank accession number MN908947) was codon-optimized for Chinese Hamster Ovary (CHO) cells and synthesized by GenScript (Piscataway, NJ, USA). Within the construct, the Spike glycoprotein was preceded by its natural N-terminal signal peptide and fused at the C-terminus to human resistin (accession number NP_001180303.1, amino acids 23-108). Mutations were added to stabilize the generated Spike protein as previously described; amino acids 682-685 (RRAR) and 986-987 (KV) were replaced with GGAS and PP, respectively ^42,43^. Constructs were then cloned into the pTT241 plasmid that did not encode C-terminal FLAG/His affinity tags. Expression constructs for VOC Spike variants were prepared by re-synthesizing and replacing restriction fragments encompassing mutations present in the Beta (SmT1v3-B) (D80A, D215G, 241del, 242del, 243del, K417N, E484K, N501Y, D614G, A701V) and Delta (SmT1v3-D) (T19R, G142D, E156-, F157-, R158G, L452R, T478K, D614G, P681R, D950N) variants, while maintaining the codon-optimized sequences of the remaining amino acids used for SmT1v3-Reference (R) expression. Stably transfected pools were established by methionine sulfoximine selection using the CHO^2353™^ cell line and used for 10-day fed-batch productions with cumate induction as described ^41^. SmT1v3-R and –D were purified using a one-step affinity method with NGL COVID-19 Spike Protein Affinity Resin (Repligen, Waltham, MA, USA). 150-250 ml of supernatant was applied to a 15-ml column by gravity flow; Dulbecco’s Phosphate Buffered Saline (DPBS) was used for column equilibration and wash steps while the elution buffer consisted of 100 mM sodium acetate, pH 3.5. SmT1v3-B was purified using a proprietary multi-step non-affinity-based process. After purification, proteins were formulated by buffer exchange (using P100 desalting columns (CentriPure, EMP Biotech, Berlin, Germany)) or tangential flow filtration (Minimate 30K cassettes (Pall, Port Washington, NY, USA)) in DPBS adjusted to pH 7.8 at protein concentrations of 0.9-1.1 mg/ml. Purified proteins analyzed by sodium dodecyl sulfate-polyacrylamide gel electrophoresis (SDS-PAGE) and analytical size-exclusion ultra-high performance liquid chromatography (SEC-UPLC). SEC-UPLC was run on an Acquity H-Class Bio UPLC system (Wyatt Technology, Santa Barbara, CA, USA) in phosphate buffered saline (PBS) + 0.02% Tween-20 on a 4.6 × 300 mm Acquity BEH450 column (2.5 μm bead size; Waters Limited, Mississauga, ON, Canada) coupled to a miniDAWN Multi-Angle Light Scattering (MALS) detector and Optilab T-rEX refractometer (Wyatt Technology). The identity and purity of the antigens was also confirmed by mass spectrometry. Absence of endotoxin contamination was verified using Endosafe cartridge-based Limulus amebocyte lysate tests (Charles River Laboratories, Charleston, SC, USA).

BDX301 was prepared from cultures of *Neisseria meningitidis* by detergent extraction and ethanol precipitation as described previously^44^. AdjuPhos^™^ (Invivogen, SanDiego, CA, USA) was prepared as per manufacturer’s instructions.

### Immunization and sample collection

Antigen and adjuvant vaccine components were diluted in DPBS (Cytiva, Marlborough, Massachusetts, USA) and then admixed in glass vials (Thermo Fisher Scientific) prior to administration. Antigen and adjuvant doses were chosen based on findings from previous studies. Mice were first anesthetized with isofluorane then immunized by intramuscular injection (100 µL split equally into the left and right tibialis anterior muscles) or by intranasal administration (10 µL per nare) on Days 0 and 21. Blood was collected from isofluorane-anesthetized mice, via the submandibular vein on Days 21 (prior to boost) and 35. All mice were humanely euthanized on Day 35 to collect bronchoalveolar lavage (BAL) fluid. Hamsters were first anesthetized with 3-5% isofluorane and then immunized by intramuscular injection (100 µL) into the left tibialis anterior muscle or by intranasal administration (20 µL per nare) on Days 0 and 21. Blood was collected from isoflurane-anesthetized hamsters, via the anterior vena cava on Days 20 and 35. On Day 42, all Hamsters were anesthetized by intraperitoneal injection with 100 mg/kg of ketamine and 7 mg/kg of xylazine prior to an intranasal challenge with 1 × 10^5^ PFU of SARS-CoV-2. Hamsters were monitored daily for body weight change and clinical signs. On Day 47, hamsters were euthanized and nasal turbinate and the left lung were collected for assessment of viral load by plaque assay as described below.

### Anti-Spike IgG ELISA

Anti-Spike total IgG titers in serum were quantified by ELISA. Briefly, 96–well high-binding ELISA plates (Thermo Fisher Scientific) were coated overnight at room temperature (RT) with 100 µL of 0.3 µg/mL Spike protein (same as used for immunization) diluted in PBS. Plates were washed five times with PBS/0.05% Tween20 (PBS-T; Sigma-Aldrich, St. Louis, MO, USA), and then blocked for 1 hour at 37 °C with 200 µL 10% fetal bovine serum (FBS; Thermo Fisher Scientific) in PBS. After the plates were washed five times with PBS-T, 3.162-fold serially diluted samples in PBS-T with 10% FBS were added in 100 µL volumes and incubated for 1 hour at 37 °C. After five washes with PBS-T (Sigma-Aldrich), 100 µL of goat anti-mouse IgG -HRP (1:4,000, Southern Biotech, Birmingham, AL, USA) or goat anti-hamster IgG-HRP (1:32,000, Southern Biotech) was added for 1 hour at 37 °C. After five washes with PBS-T, 100 µL/well of the substrate o-phenylenediamine dihydrochloride (OPD, Sigma-Aldrich) diluted in 0.05 M citrate buffer (pH 5.0) was added. Plates were developed for 30 minutes at RT in the dark. The reaction was stopped with 50 µL/well of 4N H_2_SO_4_. Bound IgG Abs were detected spectrophotometrically at 450 nm. Titers for IgG in serum were defined as the dilution that resulted in an absorbance value (OD 450) of 0.2 and were calculated using XLfit software (ID Business Solutions, Guildford, UK). Samples that do not reach the target OD were assigned the value of the lowest tested dilution (i.e. 10) for analysis purposes.

### Anti-Spike IgA ELISA

Anti-Spike IgA titers were also measured by indirect ELISA. Specifically, 96-well high-binding ELISA plates (Thermo Fisher Scientific) were coated overnight at 4 °C with 100 μL of 0.8 μg/mL (80 ng/well) of Spike protein diluted in PBS. After washing plates with PBST, wells were blocked with 200 μL 3% Skim milk (Thermo Fisher Scientific) in PBS and incubated at 37 °C for 2 hours and washed with PBST. 100 μL of sample was loaded on the plates using half-log serial dilution (equivalent to 3.162 x serial dilution starting at 1:10) and incubated at 37 °C for 1 hour. Plates were washed again with PBST and 100 μL of goat anti-mouse IgA-HRP (Abcam, Cambridge, UK) diluted 1:10,000 added to each well. Plates were incubated at 37 °C for 45 min and washed with PBST. 100 μL of TMB substrate Sure-blue reserve one-component (Mandel Scientific Company Inc., Guelph, ON, Canada) was then added to each well and reaction stopped after 10 min incubation at room temperature by adding 100 μL of TMB stop solution one-component (Mandel Scientific Company Inc., Guelph, ON, Canada). Bound IgA Abs were detected spectrophotometrically at 450 nm, and IgA antibody titers were determined as above for IgG.

### Vero E6 Cell-based Surrogate SARS-CoV-2 Neutralization Assay

This surrogate neutralization assay was performed similarly to conventional pseudoparticle neutralization assays^45^. Briefly, the ability of labeled SARS-CoV-2 Spike trimers to bind the surface of Vero E6 cells following co-incubation with sera/plasma was measured. Vero E6 cells were maintained in RPMI 1640 supplemented with 10% FBS, 1% penicillin/streptomycin, 20mM HEPES, 1x non-essential amino acids, 1x Glutamax, 50µM 2-mercaptoethanol (all from Thermo Fisher Scientific) at 37°C with 5% CO2. Soluble SmT1v3, was biotinylated and isolated from free biotin using EZ-Link™ NHS-LC-LC-Biotin (Thermo Fisher Scientific) according to manufacturer’s instructions. Indicated dilutions of mouse/hamster serum were incubated with 250 ng of biotinylated Spike and 1×10^5^ Vero E6 cells (ATCC® CRL-1586™) in the presence of PBS, 0.05% sodium azide (Thermo Fisher Scientific) and 1% bovine serum albumin (BSA; Rockland, Philadelphia, PA, USA) within a 96-well V-bottom plate (Nunc™, Thermo Fisher Scientific) for 1 hour at 4 °C, while protected from light. Regardless of serum concentration, the final volume of all samples was normalized to 150 µL. Cells were washed with PBS + 0.05% sodium azide + 1% BSA and incubated with Streptavidin-phycoerythrin conjugate (Thermo Fisher Scientific) for 1 hour at 4 °C. After another wash, the cells were fixed using CytoFix™ (Becton Dickinson, Franklin Lakes, NJ, USA) and resuspended in wash buffer + 5mM EDTA for acquisition on an LSR Fortessa™ (Becton Dickinson). Percent neutralization was calculated using the Geometric Mean Fluorescence Intensity (MFI) of PE (on singlet cell population) using FlowJo™ 10 analysis software (Becton Dickinson) as shown in the following formula: % neutralization = 100 – (100 × (Geometric MFI for PE of test sample – Geometric MFI for PE of negative control sample (i.e. cells incubated only with Streptavidin-PE and without Spike protein)) / (Geometric MFI for PE of positive control sample (i.e., cells incubated with Spike without serum/plasma) – Geometric MFI for PE of negative control sample)). For analysis purposes, samples with calculated values ≤ 0 were assigned a value of 0.

#### Plaque Assay

This assay was performed exclusively within a containment level 3 facility (CL3). Whole lung from each hamster was homogenized in 1 mL PBS. The samples were centrifuged and the clarified supernatants were used in a plaque assay. The plaque assay, in brief, was carried out by diluting the clarified lung homogenate in a 1 in 10 serial dilution in infection media (1x DMEM, high glucose media supplemented with 1x non-essential amino acid, 100 U/mL penicillin-streptomycin, 1 mM sodium pyruvate, and 0.1% bovine serum albumin). Vero E6 cells were infected for 1 h at 37°C before the inoculum was removed and overlay media was added, which consisted of infection media with 0.6% ultrapure, low-melting point agarose. The cells were incubated at 37°C/5% CO_2_ for 72 h. After incubation, cells were fixed with 10% formaldehyde and stained with crystal violet. Plaques were enumerated and PFU was determined per gram of lung tissue.

#### Plaque reduction neutralization tests (PRNT)

All steps carried out for the PRNT assay was performed in a CL3 facility. Serum samples were inactivated at 56°C for 30 min and stored on ice. A 1-in-2 serial dilution was carried out using inactivated serum. Diluted serum was incubated with equal volume containing 100 PFU of SARS-CoV-2 at 37°C for 1 h, followed by infection of Vero E6 cells. Adsorption of virus were carried out for 1 h at 37°C. Inoculum was removed after adsorption and overlay media as described above was added over the infected cells. The assay was incubated at 37°C/5% CO_2_ for 72 h. After incubation, cells were fixed with 10% formaldehyde and stained with crystal violet. Controls included naïve animal serum, as well as a no serum, virus-only back-titer control. PRNT_80_ is defined as the highest dilution of serum that results in 80% reduction of plaque-forming units. Samples that did not result in an 80% reduction in PFUs were assigned the value of the lowest tested dilution (i.e. 40) for analysis purposes.

### Statistical analysis

Data were analyzed using GraphPad Prism® version 9 (GraphPad Software, San Diego, CA, USA). Statistical significance of the difference between groups was calculated by one-way or two-way analysis of variance (ANOVA) followed by post-hoc analysis using Tukey’s (comparison across all groups) multiple comparison test. A Student’s t-test was applied when analyzing the significance of the difference between the antigen-specific IgG/IgA ratios in the BAL and serum. Data was log transformed (except for % neutralization, IgG/IgA ratio and % body weight loss) prior to statistical analysis. For all analyses, differences were considered to be not significant with p > 0.05. Significance was indicated in the graphs as follows: *p < 0.05, **p < 0.01, ***p<0.001 and ****: p<0.0001.

## Author Contributions

FCS, BA, CDZ, TRC and MH conceived and designed the studies. LD, MS, YD, GA, TMR, RD, BAH, DD, JB, SM, JZ, and MG contributed to the synthesis of the vaccine components and/or execution of the experiments. FCS, BA, LD, AT, GA, TMR, RD, BAH, DD, JB, SRM and CDZ analyzed the data. FCS, BA, JZ and CDZ took the lead in writing the manuscript. All authors provided critical feedback and helped to shape the research, data analysis and manuscript.

## Institutional Review Board Statement

Animals were maintained at the small animal facility of the National Research Council (NRC) Canada in accordance with the guidelines of the Canadian Council on Animal Care. All procedures performed on animals in this study were approved by our Institutional Review Board (NRC Human Health Therapeutics Animal Care Committee) and covered under animal use protocols 2020.06 & 2020.10. All experiments were carried out in accordance with the ARRIVE guidelines.

## Data Availability Statement

The data presented in this study are available on request from the corresponding author. The data are not publicly available due to privacy concerns.

## Acknowledgments

The authors would like to acknowledge the technical contribution of many members of the Mammalian Cell expression Section and Animal Resources Group of the NRC-HHT.

## References

1. Davis, C. et al. Reduced neutralisation of the Delta (B.1.617.2) SARS-CoV-2 variant of concern following vaccination. PLoS Pathog. 17, e1010022 (2021).

2. Pegu, A. et al. Durability of mRNA-1273 vaccine-induced antibodies against SARS-CoV-2 variants. Science 373, 1372–1377 (2021).

3. Tatsi, E.-B., Filippatos, F. & Michos, A. SARS-CoV-2 variants and effectiveness of vaccines: a review of current evidence. Epidemiol. Infect. 149, e237 (2021).

4. Baden, L. R. et al. Efficacy and Safety of the mRNA-1273 SARS-CoV-2 Vaccine. N. Engl. J. Med. 384, 403–416 (2021).

5. Walsh, E. E. et al. Safety and Immunogenicity of Two RNA-Based Covid-19 Vaccine Candidates. N. Engl. J. Med. 383, 2439–2450 (2020).

6. Folegatti, P. M. et al. Safety and immunogenicity of the ChAdOx1 nCoV-19 vaccine against SARS-CoV-2: a preliminary report of a phase 1/2, single-blind, randomised controlled trial. The Lancet 396, 467– 478 (2020).

7. Castells, M. C. & Phillips, E. J. Maintaining Safety with SARS-CoV-2 Vaccines. N. Engl. J. Med. 384, 643– 649 (2021).

8. Hatziantoniou, S., Maltezou, H. C., Tsakris, A., Poland, G. A. & Anastassopoulou, C. Anaphylactic reactions to mRNA COVID-19 vaccines: A call for further study. Vaccine 39, 2605–2607 (2021).

9. Greinacher, A. et al. Thrombotic Thrombocytopenia after ChAdOx1 nCov-19 Vaccination. N. Engl. J. Med. 384, 2092–2101 (2021).

10. Hayawi, K., Shahriar, S., Serhani, M. A., Alashwal, H. & Masud, M. M. Vaccine versus Variants (3Vs): Are the COVID-19 Vaccines Effective against the Variants? A Systematic Review. Vaccines 9, 1305 (2021).

11. Hou, Y. J. et al. SARS-CoV-2 Reverse Genetics Reveals a Variable Infection Gradient in the Respiratory Tract. Cell 182, 429-446.e14 (2020).

12. Holmgren, J. & Czerkinsky, C. Mucosal immunity and vaccines. Nat. Med. 11, S45–S53 (2005).

13. Bilsborough, J. & Viney, J. L. Gastrointestinal dendritic cells play a role in immunity, tolerance, and disease. Gastroenterology 127, 300–309 (2004).

14. Burt, D. et al. Proteosome-adjuvanted intranasal influenza vaccines: advantages, progress and future considerations. Expert Rev. Vaccines 10, 365–375 (2011).

15. Fujkuyama, Y. et al. Novel vaccine development strategies for inducing mucosal immunity. Expert Rev. Vaccines 11, 367–379 (2012).

16. Rhee, J. H., Lee, S. E. & Kim, S. Y. Mucosal vaccine adjuvants update. Clin. Exp. Vaccine Res. 1, 50–63 (2012).

17. Gupta, T. & Gupta, S. K. Potential adjuvants for the development of a SARS-CoV-2 vaccine based on experimental results from similar coronaviruses. Int. Immunopharmacol. 86, 106717 (2020).

18. Patel, G. B. & Chen, W. Archaeal lipid mucosal vaccine adjuvant and delivery system. Expert Rev. Vaccines 9, 431–440 (2010).

19. Mallett, C. P. et al. Intransal or intragastric immunization with proteosome-Shigella lipopolysaccharide vaccines protects against lethal pneumonia in a murine model of Shigella infection. Infect. Immun. 63, 2382–2386 (1995).

20. Orr, N., Robin, G., Cohen, D., Arnon, R. & Lowell, G. H. Immunogenicity and efficacy of oral or intranasal Shigella flexneri 2a and Shigella sonnei proteosome-lipopolysaccharide vaccines in animal models. Infect. Immun. 61, 2390–2395 (1993).

21. Cao, W. et al. Nasal delivery of Protollin-adjuvanted H5N1 vaccine induces enhanced systemic as well as mucosal immunity in mice. Vaccine 35, 3318–3325 (2017).

22. Jones, T. et al. A nasal Proteosome^™^ influenza vaccine containing baculovirus-derived hemagglutinin induces protective mucosal and systemic immunity. Vaccine 21, 3706–3712 (2003).

23. Lambkin-Williams, R. et al. An Intranasal Proteosome-Adjuvanted Trivalent Influenza Vaccine Is Safe, Immunogenic & Efficacious in the Human Viral Influenza Challenge Model. Serum IgG & Mucosal IgA Are Important Correlates of Protection against Illness Associated with Infection. PloS One 11, e0163089 (2016).

24. Hu, M. C. et al. Intranasal Protollin-formulated recombinant SARS S-protein elicits respiratory and serum neutralizing antibodies and protection in mice. Vaccine 25, 6334–6340 (2007).

25. Akache, B. et al. Immunogenic and efficacious SARS-CoV-2 vaccine based on resistin-trimerized spike antigen SmT1 and SLA archaeosome adjuvant. Sci. Rep. 11, 21849 (2021).

26. Arunachalam, P. S. et al. Adjuvanting a subunit COVID-19 vaccine to induce protective immunity. Nature 594, 253–258 (2021).

27. Yilmaz, I. C. et al. Development and preclinical evaluation of virus-like particle vaccine against COVID-19 infection. Allergy 77, 258–270 (2022).

28. Akache, B., Stark, F. C., Agbayani, G., Renner, T. M. & McCluskie, M. J. Adjuvants: Engineering Protective Immune Responses in Human and Veterinary Vaccines. Methods Mol. Biol. Clifton NJ 2412, 179–231 (2022).

29. Bartsch, Y. C. et al. Discrete SARS-CoV-2 antibody titers track with functional humoral stability. Nat. Commun. 12, (2021).

30. Tada, T. et al. Convalescent-Phase Sera and Vaccine-Elicited Antibodies Largely Maintain Neutralizing Titer against Global SARS-CoV-2 Variant Spikes. mBio 12, e0069621 (2021).

31. Imai, M. et al. Syrian hamsters as a small animal model for SARS-CoV-2 infection and countermeasure development. Proc. Natl. Acad. Sci. 117, 16587–16595 (2020).

32. Ella, R. et al. Efficacy, safety, and lot to lot immunogenicity of an inactivated SARS-CoV-2 vaccine (BBV152): a, double-blind, randomised, controlled phase 3 trial. medRxiv (2021).

33. King, R. G. et al. Single-Dose Intranasal Administration of AdCOVID Elicits Systemic and Mucosal Immunity against SARS-CoV-2 and Fully Protects Mice from Lethal Challenge. Vaccines 9, 881 (2021).

34. Cromer, D. et al. Neutralising antibody titres as predictors of protection against SARS-CoV-2 variants and the impact of boosting: a meta-analysis. Lancet Microbe 3, e52–e61 (2022).

35. Riemersma, K. K. et al. Shedding of Infectious SARS-CoV-2 Despite Vaccination. medRxiv (2021).

36. Langel, S. N. et al. Oral and intranasal Ad5 SARS-CoV-2 vaccines decrease disease and viral transmission in a golden hamster model. BioRxiv (2021).

37. van Doremalen, N. et al. Intranasal ChAdOx1 nCoV-19/AZD1222 vaccination reduces viral shedding after SARS-CoV-2 D614G challenge in preclinical models. Sci. Transl. Med. 13, eabh0755 (2021).

38. Lund, F. E. & Randall, T. D. Scent of a vaccine. Science 373, 397–399 (2021).

39. Tioni, M. F. et al. One mucosal administration of a live attenuated recombinant COVID-19 vaccine protects nonhuman primates from SARS-CoV-2. 2021.07.16.452733 (2021).

40. Stuible, M. et al. Optimization of a high-cell-density polyethylenimine transfection method for rapid protein production in CHO-EBNA1 cells. J. Biotechnol. 281, 39–47 (2018).

41. Isho, B. et al. Persistence of serum and saliva antibody responses to SARS-CoV-2 spike antigens in COVID-19 patients. Sci. Immunol. 5, (2020).

42. Wrapp, D. et al. Cryo-EM structure of the 2019-nCoV spike in the prefusion conformation. Science 367, 1260–1263 (2020).

43. Pallesen, J. et al. Immunogenicity and structures of a rationally designed prefusion MERS-CoV spike antigen. Proc. Natl. Acad. Sci. U. S. A. 114, E7348–E7357 (2017).

44. US Patent for Compositions and methods for activating innate and allergic immunity Patent (Patent # 9,433,672 issued September 6, 2016) - Justia Patents Search. https://patents.justia.com/patent/9433672.

45. Wrapp, D. et al. Structural Basis for Potent Neutralization of Betacoronaviruses by Single-Domain Camelid Antibodies. Cell 181, 1004-1015.e15 (2020).

